# Search party flock formations in migrating birds

**DOI:** 10.64898/2026.09.28.755100

**Authors:** Tomas Svensson, Richard Ottvall

## Abstract

From the iconic V-formations of geese to the spectacular murmurations of starlings and communal roosting of millions of bramblings, bird flocks keep sparking wonder and scientific curiosity. In science, common themes include how flocking formation impacts the energy expenditure during migration, how it affects risks of predation, how it relates to information spread on food abundance, and how birds manoeuvre in dense flocks. Here, we report on observations of a very different type of flock formation: migrating terns over open sea flying side by side in a search party manner – strikingly orthogonal to their heading – with interbird distances up towards 300 m and flocks of around ten birds covering widths up to one kilometer. Such sparse formations of migrating birds stand in great contrast to the classes of bird flocks previously described in the literature. We present simultaneous field observations and ship-based radar measurements of flock characteristics and argue that this is a form of fly-and-forage migration that may be a common, but overlooked, network foraging strategy of surface feeding birds: while most migrants cannot combine migration and foraging, the situation is different for surface feeders like terns.

## 1 Introduction

Flocking of birds have fascinated mankind in all times and steadily remains a topic of both public and scientific interest [1, 2]. The phenomenon comes with a rich flora of both spectacular and scientifically intriguing topics. The most famous example is perhaps the V-formations of geese and other species of waterfowl that – alongside bringing senses of joy of spring arrival and wonder over bird migration – remains a topic of active research in the field of energy savings via flight formation [3, 4]. Another famous example is the study of how birds manoeuvre in dense flocks, where the many detailed investigations of the murmurations of starlings can serve as a fascinating point of entry [5]. Yet another vast area of research is the study of how formation of flocks relates to food availability and predation pressure. Aggregation and flocking occur on many scales. In one end there are, for example, ocean-wide recruitment to multi-species seabird feeding hotspots in the North Atlantic [6, 7] and continent-level recruitment of landbirds such as the regular communal roosting of starlings [8], flocks of millions of wintering bramblings during beech mast years [9, 10] and mass concentrations of roosting sandhill cranes during spring migration [11]. In the other end, we have the ubiquitous occurrence of breeding colonies [12, 13], from the extreme case of millions of red-billed queleas in Africa and little auks in Greenland [14, 15] to colonies of a few pairs, small scale communal roosting [16, 17] and formation of smaller flocks for group foraging. The present work falls within the vast field of research on group foraging. A central theme in this area is the understanding of how group formation and social interaction impact foraging efficiency [18, 19]. In addition to the well-known phenomenon of dense flocks of foraging birds – a form of gregariousness that e.g. reduce predation risk and vigilance [20, 21, 22] – there is also the concept of network foraging. Network foraging is a form of local enhancement that concerns sparse groups of individuals in visual contact that search for food independently but reacts to the success of its neighbours and aggregate when food is found. This foraging behaviour is observed in many seabirds, vultures and corvids [23, 24, 25, 13]. As an example, GPS loggers have revealed that cape gannets, a fish-eating seabird, are recruited to feeding aggregations at distances up to 40 km [26]. There are even indications that there is a link between the white plumage of many seabirds and social foraging [27], as well as reports that fishing success increase with flock size so that even the bird that found the food resource can benefit from network foraging [28].

Here, we report on a type of network foraging that stand in great contrast to the classical examples. During combined human observation and radar monitoring conducted from a survey ship 40 km offshore in the Baltic Sea it was discovered that migrating flocks of terns flew in sparse search party formations with birds spread out orthogonally to their migration heading. Such a flock structure differs significantly from both the staggered line formations (V- and echelon formations) exhibited by larger birds like waterfowls, pelicans, cranes, ibises, gulls and some waders, and from the cluster formations that for example dominates among smaller birds (cf. [1, 29]). We present the combined analysis of collected radar data and field observations, reporting interbird separations up to almost 300 m and groups of around ten terns spanning widths up to 1 km. We argue that this is both a case of fly-and-forage migration (cf. [30, 31, 32]) and an example of organized network foraging based on visual contact. Previous reports of organized group foraging includes prey herding by lines of swimming pelicans and walking waders [33, 34, 35], synchronized diving in penguins (McInnes et al., 2017) as well as passively collected radar data showing groups of unidentified foraging seabirds appearing to be “raking the ocean surface” [36].

## 2 Materials and Methods

The data used in this study was collected during a ship-based ornithological survey being part of the environmental impact assessment of a potential wind farm development between the Swedish islands of Öland and Gotland in the Baltic Sea. As shown in Figure 1, the survey ship (M/V Skoven) was anchored offshore at 56.7023°N 17.5496°E (around 40 km from both islands) during the period 16–20 May 2024.

**Figure 1.**
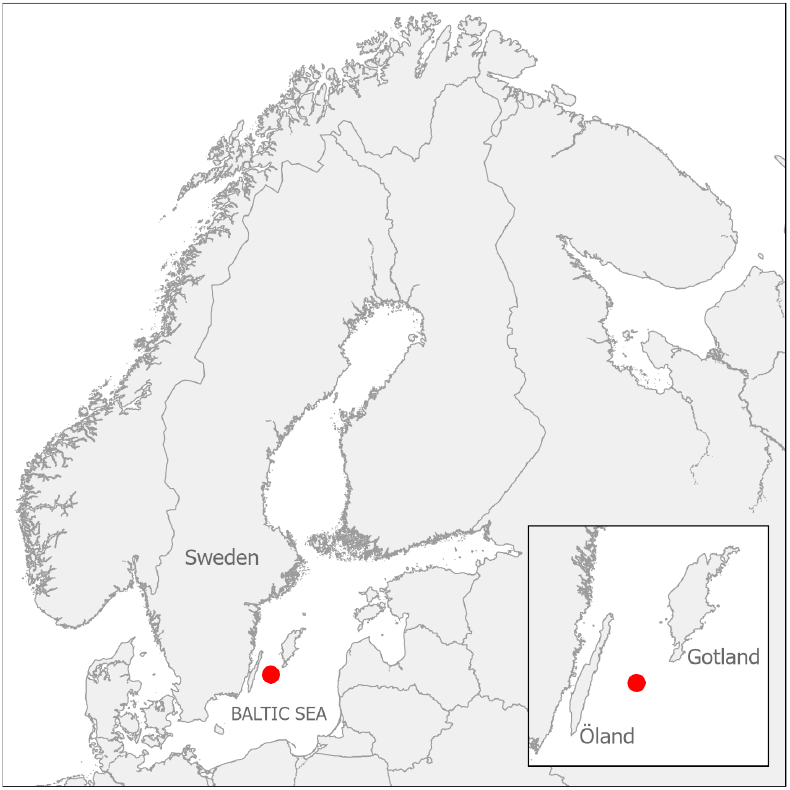
Survey location (red dot). Field observations and radar data was collected from a ship located 40 km offshore in between the two largest islands of Sweden (Öland and Gotland) during the period May 16 – May 20 (2024).

Radar data from an X-band marine radar Furuno DRS25ANX running in BirdMode was collected continuously (30 frames per minute). Field observers monitored movements along two directions from the ship using handheld binoculars as well as scopes on tripods (west and south, respectively). The two directions are marked by buoys placed at one and two kilometres from the ship. The buoys are visible in the radar images as static echoes towards the west and south. All birds seen passing between the ship and the outer buoys are registered by the observers. Birds discovered via the radar screen and then also seen visually, even if not crossing the monitored directions within 2 km, are also registered in the field notes. In additions, birds beyond 2 km and not visible in radar images are also registered if time allows. All tern flocks that were registered in the field notes and are seen in the radar images have been included in the analysis. As bird tracking with marine radars are limited to relatively calm sea conditions, the flocks analyzed were all moving in low wind speed (sea clutter, that in the context of bird monitoring acts as a radar noise floor, increases with wind and wave height and degrades the possibility to detect birds). The first step of the data analysis is manual marking of echoes from birds in the tern flocks. Due to the high spatial resolution of the radar, marking of individuals terns are often possible. However, echoes from birds flying close together overlap and cannot always be resolved. Since radar cross section of birds varies heavily with the exact orientation of the bird and the exact wing positions, echo strengths will also vary significantly from one radar images to the next. It is therefore rather common that birds can be clearly seen in one radar image to become invisible in the next. In addition, due to obstruction from objects around the radar installation, the radar quality varies with the exact azimuthal direction. These complications are however not very cumbersome as the flocks studied here are typically tracked for several minutes. The large number of radar images for each flock will provide good information on the flock formation, and there are typically several frames in which most of the birds in the flock can be seen. The agreement between the flock count as registered in the field notes is generally good, but in most cases, it is not possible to reach the same flock count from only the radar images. This is related to that there are some birds that fly too close to create separated echoes. There were, however, a couple of few cases in which the radar images showed that the field observers missed a bird or two (the spatial resolution of the radar is best in the close range, and for flocks passing within 1–2 km it is often possible to see individual birds clearly over multiple subsequent images) The manual marking of bird echoes results in datasets from which various flock parameters can be derived. Flock spread is evaluated by calculating the largest distance between birds in the flock, and the lateral width by calculating the spread perpendicular to the heading of the flock. As it is common that echoes are not seen in all frames, as described above, this analysis is done based on a number of subsequent images rather than only one image at a time. For each radar image, the lateral width is calculated based on five images (the current along with two preceding and two subsequent). In contrast, to avoid being affected by the movement along the flight direction (around 10 m/s), spread is based on only three subsequent images at a time. Since most of the flocks exhibited a pronounced search party formation (birds distributed orthogonally around the flock heading), the lateral width is often very close to the overall spread. In cases where a flocks made a nonnegligible heading change e.g. due to impact from the ship and/or buoys, the dataset is split in subsets in which the flock can be assessed as flying straight. These subsets are analysed independently in terms lateral width, heading and flight speed.

## 3 Results

In total, 34 groups of migrating terns were registered during the survey period May 16–20 (2 on May 16, 3 on May 17, 1 on May 18, 24 on May 19 and 4 on May 20). Out of these, 25 flocks were found in the radar data. Note that all flocks on May 19 could be found (thanks to calm weather), while the less calm sea conditions on the other days resulted in only one flock being found. Most of the flocks was registered as Common/Arctic Terns Sterna hirundo/paradisaea (certain species assignment often not possible due to long distances), but the analysis includes also two flocks of Black Terns Chlidonias niger, one mixed flock (one Black Tern coming together with a Common/Arctic) and one flock of Common Terns (three if including flocks not found in the radar data). The number of birds in the flocks ranged from two up to 14. In total, 15 of the 25 flocks made nonnegligible heading change, often related to the passage of the buoys and/or the ship. Flights speeds are in good agreement with previously published values for terns, black terns moving slower than the Sterna-terns [37, 38, 39, 40]. Ground speeds for all flocks are listed in the supplementary material below. The radar data show signs of potential brief stops for feeding in four of the focks. For example, one group of black terns made a ten minute stop before continuing. During this time the birds we were lost both visually and in the radar data and may thus have landed (due to the distance this could not be confirmed visually, but when they continued their journey they were again seen both visually and in radar data).

All flocks exhibited significant lateral widths, i.e. birds being distributed over significant a length perpendicular to the migration heading. Detailed data for all flocks are given as Supplementary Information (Table 1). Figure 2 shows an example of a flock exhibiting a 1km lateral width while migrating steadily towards NE (slight adjustment to southernmost buoy) and Figure 3 exemplifies the case where a flock passes closer to the ship and adjust its heading in this process. Interestingly, in the second example, the terns increased their speed of migration when passing the ship from around 10 m/s to 14 m/s. In addition, the flock formation goes from a search party to swarm and back again in this process, showing that the interaction with the ship briefly breaks the search party organization. Here, it is worth noting that it is known that birds, including terns, searching for food reduce their speed [41]. Figure 4 summarizes flock characteristics by showing how the lateral flock width and average spacing between birds varies with number of birds involved.

**Table 1:**
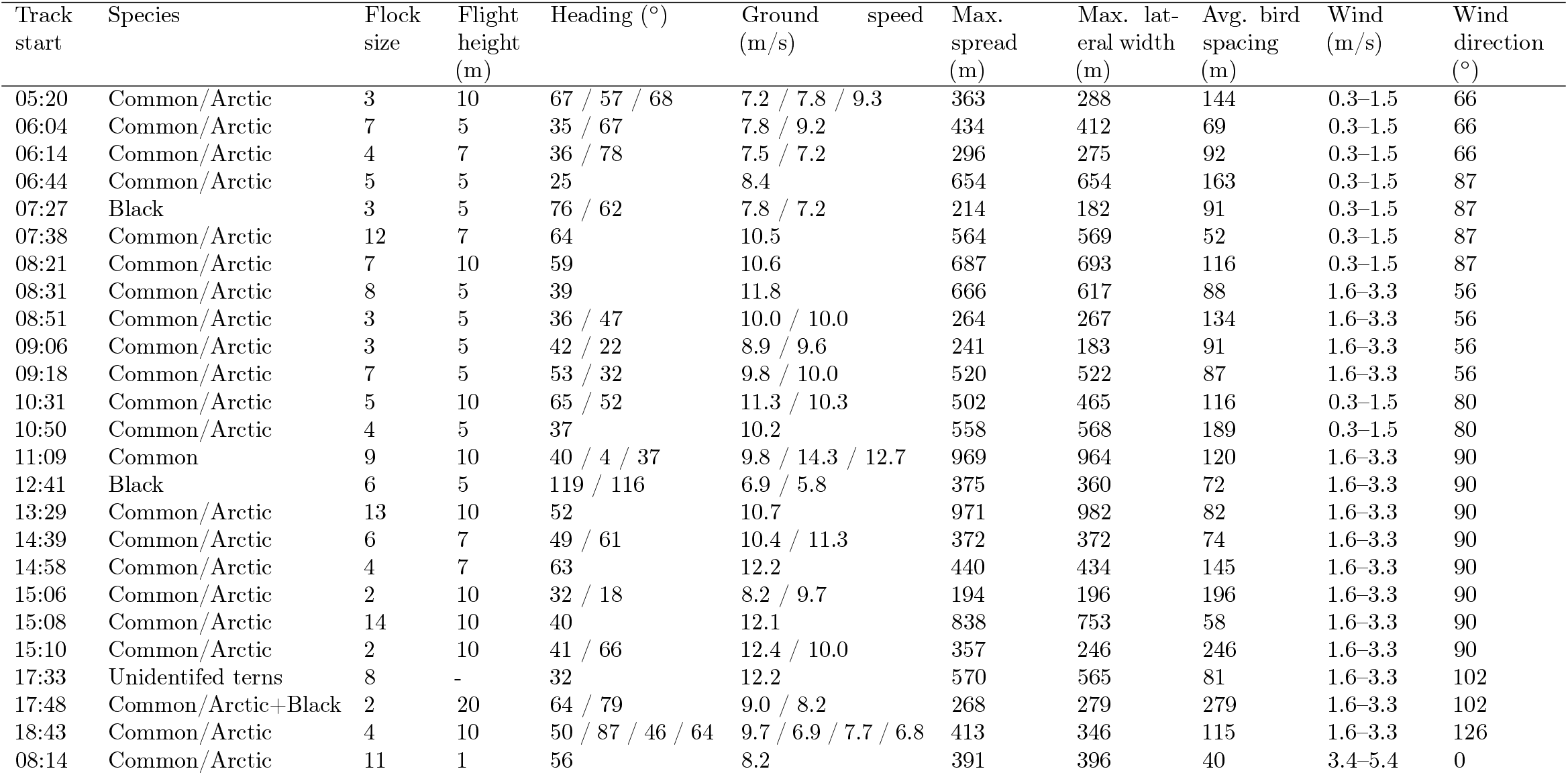
Characteristics of all flocks in this study, see Methods and Materials for details. Note that flight heights are estimated manually by field observers (not measured), Heading is in degrees clockwise from north, e.g. 90° corresponding to a flock heading towards E. Wind direction is also given in degrees clockwise from north, but here 90° means that the winds comes from E. In cases the flock made significant heading changes, heading and ground speed is presented for the al l segments (values separated by a forward slash). Note also all flocks but the last was registered on May 19 (the last flock is from May 20).

| Track start | Species | Flock size | Flight height (m) | Heading (°) | Ground speed (m/s) | Max. spread (m) | Max. lateral width (m) | Avg. bird spacing (m) | Wind (m/s) | Wind direction (°) |
| --- | --- | --- | --- | --- | --- | --- | --- | --- | --- | --- |
| 05:20 | Common/Arctic | 3 | 10 | 67 / 57 / 68 | 7.2 / 7.8 / 9.3 | 363 | 288 | 144 | 0.3–1.5 | 66 |
| 06:04 | Common/Arctic | 7 | 5 | 35 / 67 | 7.8 / 9.2 | 434 | 412 | 69 | 0.3–1.5 | 66 |
| 06:14 | Common/Arctic | 4 | 7 | 36 / 78 | 7.5 / 7.2 | 296 | 275 | 92 | 0.3–1.5 | 66 |
| 06:44 | Common/Arctic | 5 | 5 | 25 | 8.4 | 654 | 654 | 163 | 0.3–1.5 | 87 |
| 07:27 | Black | 3 | 5 | 76 / 62 | 7.8 / 7.2 | 214 | 182 | 91 | 0.3–1.5 | 87 |
| 07:38 | Common/Arctic | 12 | 7 | 64 | 10.5 | 564 | 569 | 52 | 0.3–1.5 | 87 |
| 08:21 | Common/Arctic | 7 | 10 | 59 | 10.6 | 687 | 693 | 116 | 0.3–1.5 | 87 |
| 08:31 | Common/Arctic | 8 | 5 | 39 | 11.8 | 666 | 617 | 88 | 1.6–3.3 | 56 |
| 08:51 | Common/Arctic | 3 | 5 | 36 / 47 | 10.0 / 10.0 | 264 | 267 | 134 | 1.6–3.3 | 56 |
| 09:06 | Common/Arctic | 3 | 5 | 42 / 22 | 8.9 / 9.6 | 241 | 183 | 91 | 1.6–3.3 | 56 |
| 09:18 | Common/Arctic | 7 | 5 | 53 / 32 | 9.8 / 10.0 | 520 | 522 | 87 | 1.6–3.3 | 56 |
| 10:31 | Common/Arctic | 5 | 10 | 65 / 52 | 11.3 / 10.3 | 502 | 465 | 116 | 0.3–1.5 | 80 |
| 10:50 | Common/Arctic | 4 | 5 | 37 | 10.2 | 558 | 568 | 189 | 0.3–1.5 | 80 |
| 11:09 | Common | 9 | 10 | 40 / 4 / 37 | 9.8 / 14.3 / 12.7 | 969 | 964 | 120 | 1.6–3.3 | 90 |
| 12:41 | Black | 6 | 5 | 119 / 116 | 6.9 / 5.8 | 375 | 360 | 72 | 1.6–3.3 | 90 |
| 13:29 | Common/Arctic | 13 | 10 | 52 | 10.7 | 971 | 982 | 82 | 1.6–3.3 | 90 |
| 14:39 | Common/Arctic | 6 | 7 | 49 / 61 | 10.4 / 11.3 | 372 | 372 | 74 | 1.6–3.3 | 90 |
| 14:58 | Common/Arctic | 4 | 7 | 63 | 12.2 | 440 | 434 | 145 | 1.6–3.3 | 90 |
| 15:06 | Common/Arctic | 2 | 10 | 32 / 18 | 8.2 / 9.7 | 194 | 196 | 196 | 1.6–3.3 | 90 |
| 15:08 | Common/Arctic | 14 | 10 | 40 | 12.1 | 838 | 753 | 58 | 1.6–3.3 | 90 |
| 15:10 | Common/Arctic | 2 | 10 | 41 / 66 | 12.4 / 10.0 | 357 | 246 | 246 | 1.6–3.3 | 90 |
| 17:33 | Unidentified terns | 8 | - | 32 | 12.2 | 570 | 565 | 81 | 1.6–3.3 | 102 |
| 17:48 | Common/Arctic+Black | 2 | 20 | 64 / 79 | 9.0 / 8.2 | 268 | 279 | 279 | 1.6–3.3 | 102 |
| 18:43 | Common/Arctic | 4 | 10 | 50 / 87 / 46 / 64 | 9.7 / 6.9 / 7.7 / 6.8 | 413 | 346 | 115 | 1.6–3.3 | 126 |
| 08:14 | Common/Arctic | 11 | 1 | 56 | 8.2 | 391 | 396 | 40 | 3.4–5.4 | 0 |

**Figure 2.**
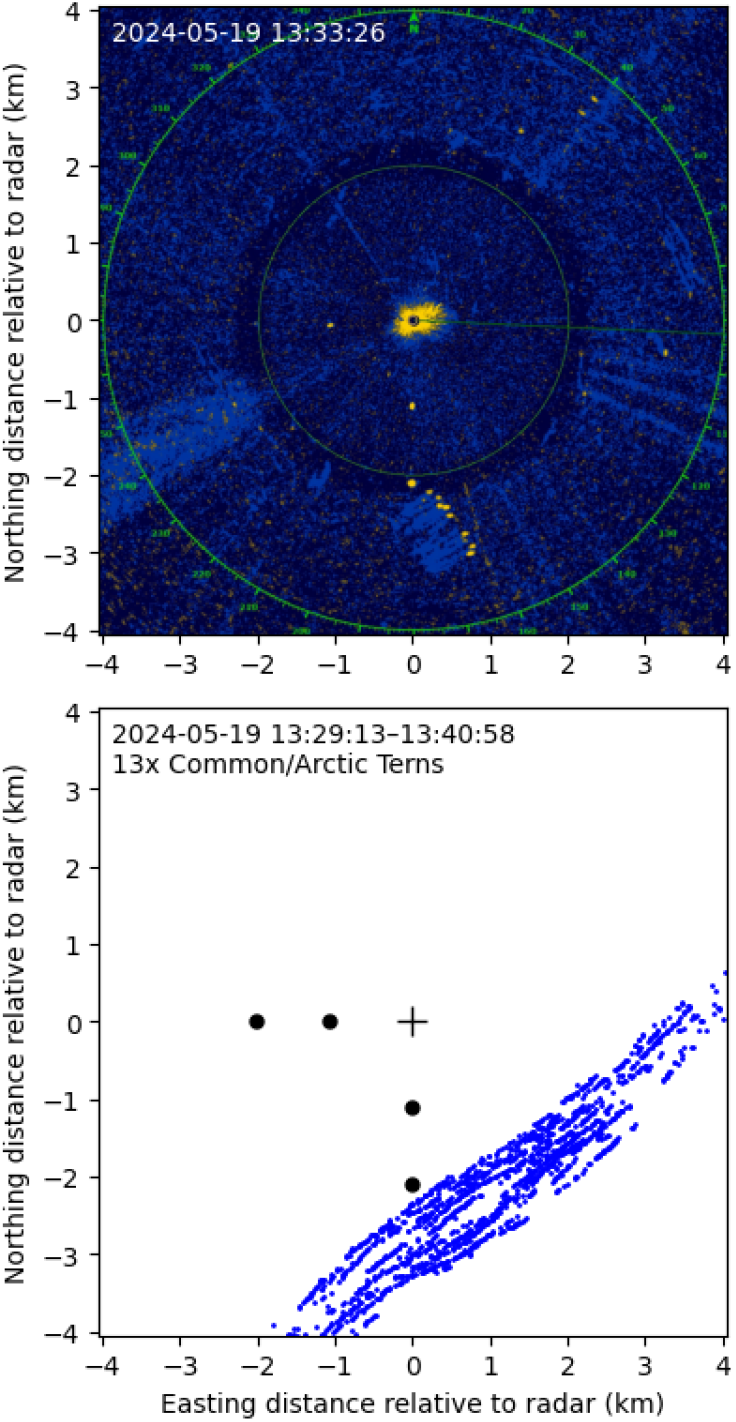
Radar screen example (top) and all individual radar echoes (bottom) for a flock of 13 Common/Arctic Terns migrating northeast while distributed side-by-side in an up to ~ 980 m wide search party formation. The flock passed just south of the 2 km buoy that marks the southwards observation direction and was tracked for above ten minutes (~ 7 km), steadily migrating northeast. In the radar image (top panel), the flock is seen as a band of yellow dots followed by radar trails in light blue (indicating old positions of moving echoes). In the middle, just around the radar, there is a region dominated by sea clutter (seen as a yellow blob). The buoys that mark the direction of manual observation are seen as static echoes (small yellow blobs) at one and two kilometres from the radar, two towards the south and two towards the west (the westernmost barely seen in this case). In the bottom panel, individual radar echoes are shown as blue dots, buoy locations as solid black dots, and the radar (ship) as a plus sign.

**Figure 3.**
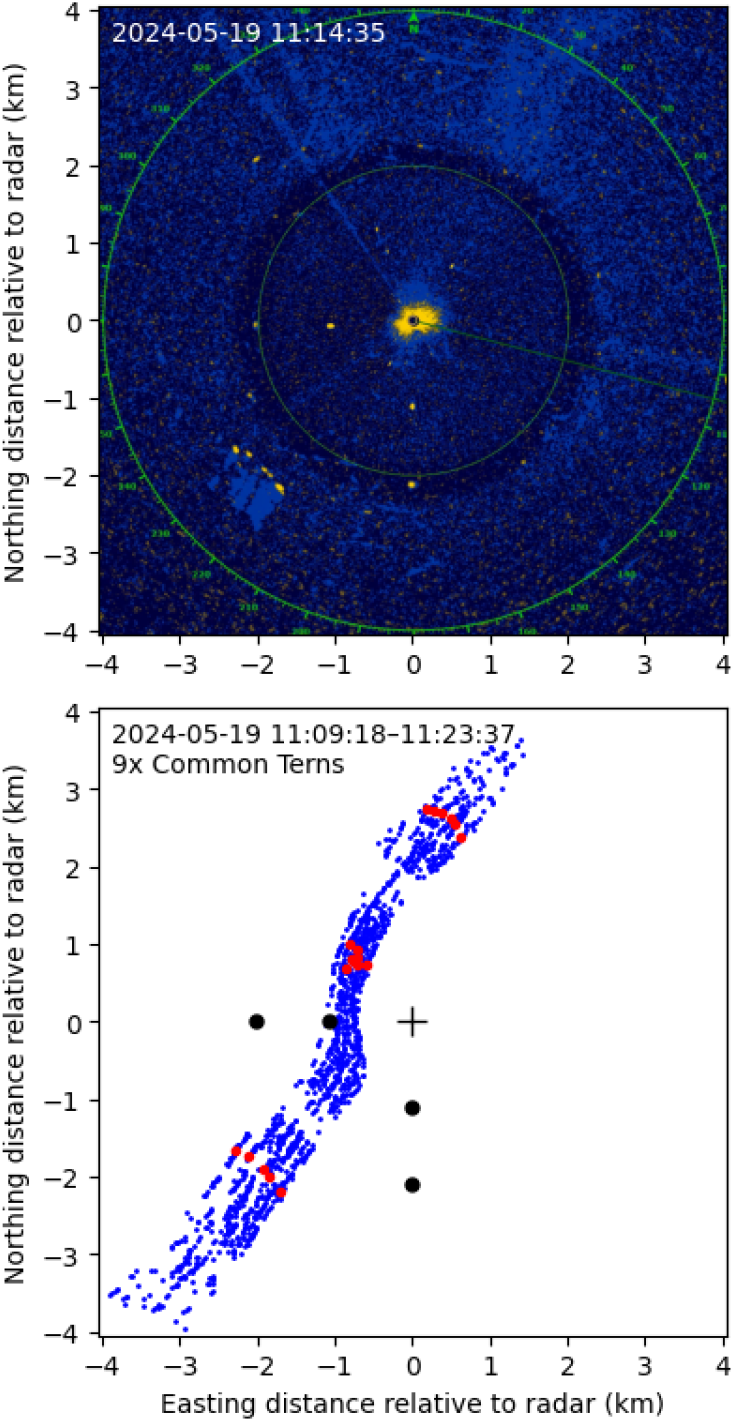
Radar screen example (top) and all individual radar echoes (bottom) for a flock of nine Common Terns that adjusted their heading when passing the ship. For information on how to interpret panels, see Figure 2. After initially having a width of ~ 930 m, the flock comes together when passing the ship/buoys, but broaden again and appear to retake its original heading soon after. This process is illustrated by showing the flock structure before, during and after the passage (red dots, echoes from three different radar frames). Note that the terns also increased their speed when passing, from 10 to 14 m/s.

**Figure 4.**
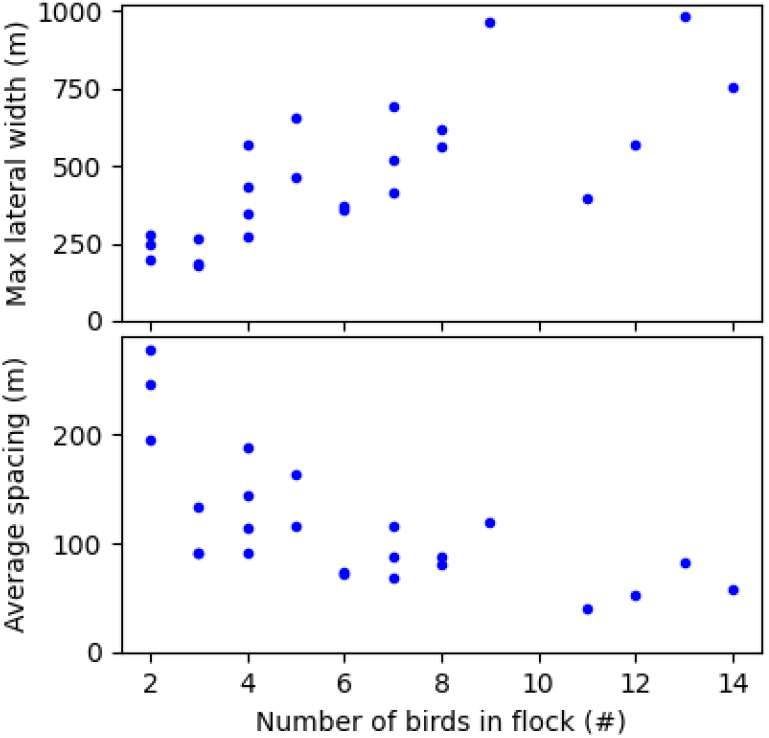
The lateral width of flocks in this study increased from around 250 meters for groups of only two birds, up to around 1000 meters for flocks around ten birds (top panel). This means that the average spacing between birds decreased with the number of birds in the flock, from above 200 meters or more for flocks of two, down to below 100 m for the larger flocks (bottom panel). Data from Table 1.

## 4 Discussion

As the observations are made offshore over 40 km from land and concerns groups of birds tracked over several kilometers while heading in expected migration directions, the terns in this study can safely be regarded as migrating rather than engaged in local foraging. In this region, the migration of common and arctic terns typically peak around the April/May shift, but migratory movements towards the northeast can be registered also well into the summer period. This is related to the later arrival of terns of northern populations, which partly pass over the Baltic [42, 43], in combination with the general delayed arrival of inexperienced breeders. For example, an extensive transponder study of common terns breeding in Germany showed that older birds (5 years and older) typically arrived during the second half of April while younger birds arrived later, with some two year olds arriving as late as in the beginning of August [44]. The migration of black terns peaks in mid May, i.e. somewhat later [45, 46]. Part of the black terns seen moving in southern Sweden may belong to the strong populations found in eastern Europe (note that one of the two flocks of black terns in this study was seen heading southeast). Since it is beyond reasonable doubt that the observed terns indeed were migrating, we argue that our observations is a case of fly-and-forage migration and, at the same time, an example of network foraging based on visual contact. The terns organize themselves as a search party hoping that patches of food will not be small enough to slip through, or with the words of Assali et al. [36], the flocks appear to be “raking the ocean surface”. In this context, a very interesting parallel is the observation that white storks fly in a manner that indicates that they search for thermals by forming similar search party formations as described here, covering lateral widths of 200–300 m, albeit with rather small inter-bird distance [47]. It is also interesting to note that since all of our observations involved terns flying low (most flocks estimated to fly 5 – 10 m above the sea surface), it seems unreasonable to believe the terns are efficiently scanning the whole lateral flock width. Instead, it appears as if the efficacy of the approach relies on that patches of food are often large enough to be comparable to the interbird spacing (i.e. the rake being dense enough despite interbird spacings of around 100 m). It should at the same time be noted that the flock structure does not form a perfect rake. The bird spacing is not perfectly equidistant, and radar echoes indicate that two or more individuals sometimes can fly close together. Limited spatial resolution in combination with large variation in radar cross sections of birds makes a more detailed analysis of flock structure difficult. Here, we instead reported on the overall lateral width and average spacing between birds. Going forward, more detailed investigations of flock structure would be valuable. Another open question is whether the search party formations are mainly seen during calm weather due to that a rougher sea surface renders fly-and-forage less fruitful Assuming that the sizes of food patches are small compared to the lateral flock width or the interbird spacing (the nearest neighbour distance), it is evident that flying in a search party flock formation can drastically improve the rate at which food abundance is found. By joining a flock of ten birds, the rate of finding small food patches may increase with a factor of ten. It is not within the scope here to review the literature of food patchiness (cf. [48]), but there are data showing that, for example, the size of fish schools appear to be on a level where search party flock formation would indeed make sense [49, 50, 51].

## Acknowledgement

We are indebted to Thomas Alerstam for insightful discussions on the topic and for encouraging us to publish these findings. We also thank Mikael Haraldsson for serving as co-observer on this survey, David Erterius (Ottvall Consulting) for his contributions to survey planning and the Skoven staff for hosting us and ensuring our safety. We also acknowledge the support from OX2.

## A Supplementary Information

